# Quantitative insights into the efficacy of genome-resolved surveys of microbial communities through ribosomal protein phylogeography and EcoPhylo

**DOI:** 10.1101/2025.01.15.633187

**Authors:** Matthew S. Schechter, Florian Trigodet, Iva A. Veseli, Samuel E. Miller, Matthew L. Klein, Metehan Sever, Loïs Maignien, Tom O. Delmont, Samuel H. Light, A. Murat Eren

## Abstract

The increasing availability of microbial genomes is essential to gain insights into microbial ecology and evolution that can propel biotechnological and biomedical advances. Recent advances in genome recovery have significantly expanded the catalogue of microbial genomes from diverse habitats. However, the ability to explain how well a set of genomes account for the diversity in a given environment remains challenging for individual studies or biome-specific databases. Here we present EcoPhylo, a computational workflow to characterize the phylogeography of any gene family through integrated analyses of genomes and metagenomes, and apply this approach to ribosomal proteins to quantify phylogeny-aware genome recovery rates in two genome-resolved investigations of the human gut and oral cavity. Our results demonstrate that EcoPhylo reveals highly resolved, reference-free, multi-domain phylogenies in conjunction with distribution patterns of individual clades across environments, providing a means to assess genome recovery in individual studies and benchmark genome collections.

## Introduction

Establishing comprehensive genome catalogues is a fundamental objective in microbiology as genomes are essential to develop insights into microbial life and to advance biotechnology and biomedicine (Eren and Banfield 2024). Indeed, the rapidly increasing number of microbial genomes (1) provides an evolutionary framework to resolve the branches of the Tree of Life (Brown et al. 2015; Spang et al. 2015), (2) enables hypothesis generation and testing through comparative genomics (Paoli et al. 2022; Al-Shayeb et al. 2022; Durrant et al. 2023; J. Chen et al. 2024a), (3) offers resources to search for novel biosynthetic capabilities and natural products (Paoli et al. 2022; J. Chen et al. 2024b), (4) contributes to the body of nucleotide data used to train biological language models (Cornman et al. 2024; Nguyen et al. 2024; Hwang et al. 2024) and more, while well-structured databases aim to consolidate and give access to the outcomes of genome recovery efforts (Parks et al. 2022; Schmidt et al. 2024).

Increasing availability of microbial genomes is a result of multiple complementary breakthroughs that include (1) advances in high-throughput or targeted cultivation that enable the recovery of isolate genomes (Jiang et al. 2016; Watterson et al. 2020; Cross et al. 2019), (2) the use of environmental shotgun sequencing that enables the recovery of metagenome-assembled genomes (MAGs) (L.-X. Chen et al. 2020), and (3) the use of microfluidics and cell sorting that enables the recovery of single amplified genomes (SAGs) (Woyke, Doud, and Schulz 2017). These strategies have not only been used in large-scale characterization of many of the Earth’s biomes (Pasolli et al. 2019; Parks et al. 2017; Pachiadaki et al. 2019; Ma et al. 2023), but also have been applied to many specific questions or niche systems that span a wide range of research priorities, collectively resulting in over 500,000 non-redundant bacterial and archaeal genomes (Parks et al. 2022). The recovery of microbial genomes is now a relatively well-established practice, yet it is not straightforward to assess (1) how taxonomic or biome-specific biases impact on genome recovery efforts, and (2) the ecological or evolutionary importance of unrecovered populations. As a result, individual studies that recover genomes, or efforts that curate biome-specific or global genomic collections, rarely offer quantitative insights into one of the key questions they aim to address: “how well do these genomes represent this environment?”.

Attempts to benchmark genome recovery often rely upon metagenomic read recruitment statistics to quantify the fraction of reads that map to genomes with the assumption that the proportion of reads recruited by a genomic collection is a proxy for the degree to which a genome collection represents the genomic fragments found in a given environment. In individual studies that reconstruct genomes directly from environmental metagenomes, the proportion of metagenomic reads that are recruited by resulting MAGs can vary from as low as 7% in the surface ocean (Delmont et al. 2018) to as high as 80% in the human gut (Carter et al. 2023). While read recruitment statistics are easy to generate and communicate, they fail to contextualize what is present in the unmapped fraction and thus leave considerable ambiguity about the microbial community. For instance, a large fraction of metagenomic reads not mapping to the genome catalogue could belong to a single organism or multiple taxonomically diverse microbes with critical ecological roles in the system. Furthermore, genome collections often systematically underrepresent certain portions of the tree of life, as the rate of genome recovery differs across taxa as a function of genome recovery methodology: while cultivation efforts often struggle to capture slow-growing organisms (Imachi et al. 2020) or those that depend on others for survival (He et al. 2015), genome-resolved metagenomics often struggle to reconstruct genomes from taxa that form highly complex populations (Giovannoni 2017; Pachiadaki et al. 2019). Altogether, biological and non-biological factors confound accurate interpretations of read recruitment results, and the ability to measure genome recovery rates requires alternative strategies that can contextualize the ecological and evolutionary relationships of organisms recovered in genome collections with environmental populations accessible through metagenomics.

One approach to gaining insight into microbial life underrepresented in genome catalogues involves the use of marker genes in metagenomic assemblies. *De novo* assembly, in which individual sequencing reads are stitched together to recover much longer contiguous segments of DNA (contigs), is common to the vast majority of genome recovery efforts. While in most cases contigs only represent fragments of genomes, they still explain a much greater genomic context than unassembled reads and give access to entire open reading frames, including phylogenetically informative marker genes. Employing such phylogenetically informative genes assembled from metagenomes enables both broad surveys of taxonomic diversity and fine-scale phylogenetic analyses. For example, RNA and DNA polymerase genes have been used to resolve viral clades and guide targeted genome recovery (Weinheimer and Aylward 2020; Gaïa et al. 2023). Additionally, marker genes have also been used to estimate the proportion of phylogenetic diversity contributed by genomes and isolate genomes in metagenomic assemblies (D. Wu et al. 2025). Furthermore, when combined with metagenomic read recruitment, such marker genes also reveal the biogeography of individual taxa, as demonstrated by studies leveraging the rpoC1 gene to uncover phylogeographic patterns in marine bacteria (Kent et al. 2019; Ustick, Larkin, and Martiny 2023).

Among all phylogenetically informative genes, ribosomal proteins represent a special class as they (1) occur as a single-copy gene in genomes across the tree of life, (2) are consistently assembled even for complex or relatively rare populations in metagenomes due to their relatively short length, and (3) contain enough phylogenetic information to delineate distinct branches of life at relatively high levels of resolution (Olm et al. 2020). Recognizing their utility, many studies have leveraged individual ribosomal proteins to analyze community composition (M. Wu and Eisen 2008; Crits-Christoph et al. 2022), integrating ribosomal protein phylogenies with metagenomic read recruitment to track individual clades of microbes (Hug et al. 2013; Emerson et al. 2016; Hamilton et al. 2016; Diamond et al. 2019; Matheus Carnevali et al. 2021). Ribosomal proteins are thus ideally suited gene markers for tracking microbial populations underrepresented within genome collections.

Here we present EcoPhylo, a workflow to simultaneously visualize the phylogenetic relationships and biogeographical distribution patterns of sequences that match any given gene family from genomes and metagenomes, and demonstrate its application to the phylogeography of ribosomal proteins for quantification of genome recovery rates across biomes. Our results show that bringing together multi-domain ribosomal protein phylogenies with distribution patterns of individual clades across environments in a single interface offers a valuable data analysis and visualization strategy to benchmark genome recovery efforts scaling from individual projects to global surveys of large genome collections and metagenomes.

## Results

### EcoPhylo enables integrated surveys of gene family phylogeography

EcoPhylo implements a computational workflow to integrate the phylogeny and biogeography of any given gene family and enables its users to track the distribution patterns and evolutionary relationships between homologous genes across environments and/or experimental conditions (Figure 1, also see Materials and Methods).

**Figure 1:**
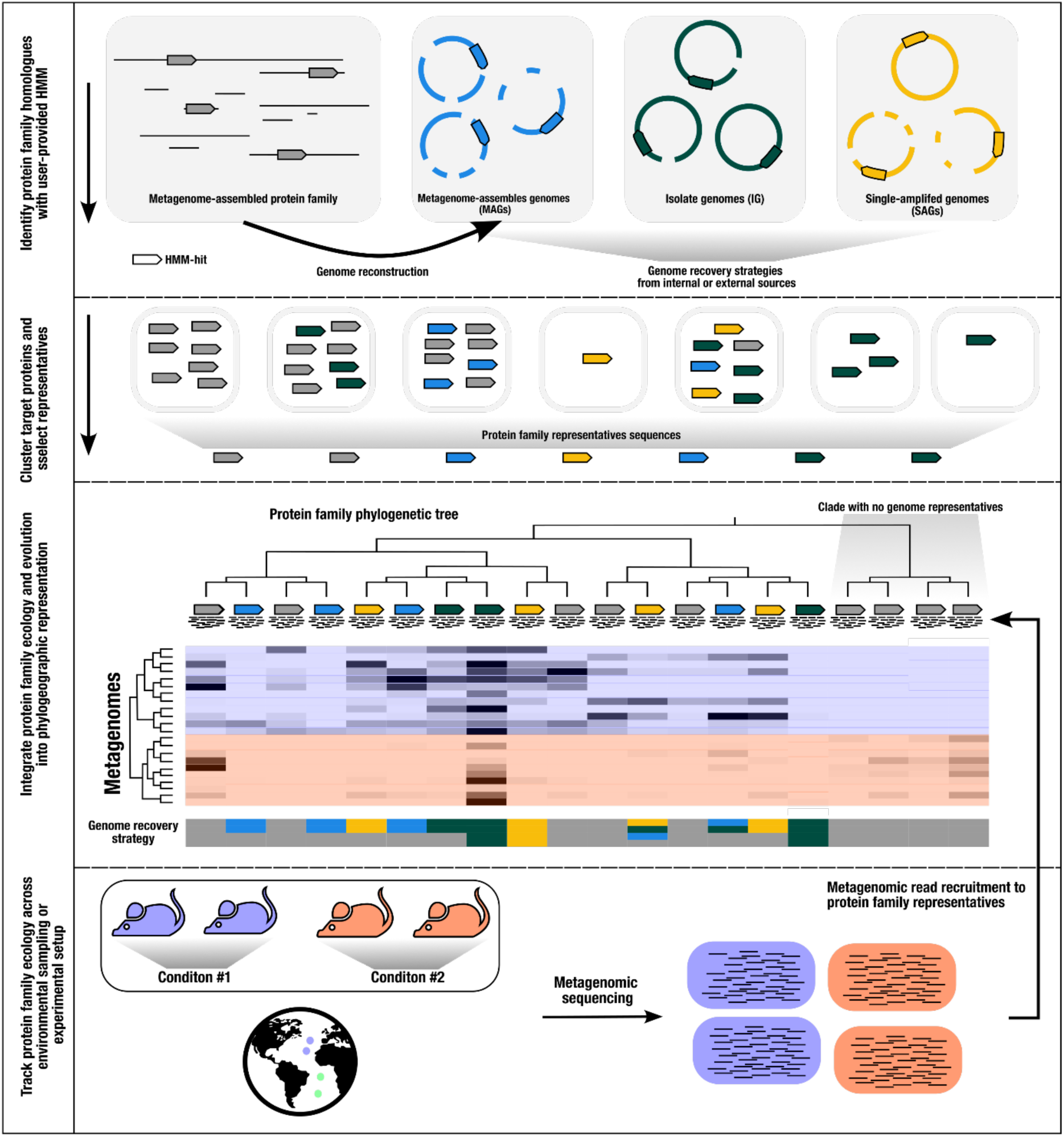
Schematic of the EcoPhylo workflow applied to a single gene family. The proposed workflow integrates biogeography from metagenomic read recruitment and protein phylogenetics to display the phylogeographical distribution of closely related lineages. When including genome sources, the workflow highlights which genome recovery strategies are more effective for sampling specific taxa. Although this manuscript focuses on ribosomal proteins, the proposed workflow is generalizable to any gene family.

When applied to phylogenetically tractable single-copy core genes, such as ribosomal proteins, in tandem with metagenomes and a genome collection, EcoPhylo identifies populations assembled in metagenomes but absent in the genomic collections (and *vice versa*), highlighting the ecological and evolutionary relevance of organisms detected through metagenomic assemblies but lacking genomic representation (Figure 1). This allows for the quantification of genome recovery rates of different methods (e.g., isolate genomes, MAGs, SAGs) across taxa and provides a means to investigate phylogenetic and ecological features of organisms without genomic representation. Importantly, the unbroken link between genes and contigs enables downstream targeted binning efforts when necessary.

Using ribosomal proteins to *de novo* characterize the phylogenetic makeup of microbiomes and benchmark genome recovery rates has numerous advantages. However, these advantages also pose noteworthy challenges. Ribosomal proteins are short protein sequences (∼300 amino acids), which substantially limits their ability to resolve deep phylogenetic branching patterns. Furthermore, their evolution is subject to strong purifying selection, as a result, the average nucleotide identity (ANI) threshold often used to define ‘species’ boundaries between whole genomes is 95% (Jain et al. 2018) increases to 99% for ribosomal protein sequences (Olm et al. 2020). Therefore, ribosomal proteins are more vulnerable than other genes to non-specific read-recruitment from closely related proteins within metagenomes. To identify criteria for reliably resolving taxa, we started our investigation by developing a series of benchmarks to optimize the use of ribosomal proteins in EcoPhylo with appropriate parameters to maximize the ecological and evolutionary signal they can offer while minimizing non-specific read recruitment. These benchmarks, which are detailed in the Supplementary information (1) inspected hidden Markov model (HMM) alignment coverage thresholds to accurately detect ribosomal proteins in genomes and metagenomes; (2) examined the copy number distribution of ribosomal protein HMMs across archaeal and bacterial genomes to only consider single-copy candidates; and (3) explored nucleotide similarity thresholds to cluster ribosomal gene sequences to maximize the taxonomic resolution of representative sequences while maintaining sufficient nucleotide distance between distinct representative sequences to reduce non-specific read recruitment from metagenomes (Supplementary information).

Based on these considerations, we implemented routines and adjusted default EcoPhylo parameters to (1) use a minimum of 80% model coverage for ribosomal protein HMMs for a match; (2) filter for complete open reading frame sequences to remove assembly artifacts; and (3) cluster HMM hits with target coverage to ensure grouping of extended open reading frames and leverage 97% nucleotide similarity as the most appropriate clustering threshold to minimize non-specific read recruitment (Supplementary information). We also compared broad ecological insights recovered from EcoPhylo to state-of-the-art taxonomic profiling tools, confirming that this framework offered qualitatively comparable results (Supplementary information). Altogether, these evaluation and optimization steps yielded EcoPhylo default parameters to obtain representative ribosomal protein sequences that are suitable for investigations of the phylogeny, biogeography, and genome recovery of populations they describe. To demonstrate the utility of the EcoPhylo workflow, we performed two case studies exploring microbial ecology and genome recovery across individual studies from the human oral cavity (Shaiber et al. 2020) and gut (Carter et al. 2023).

### Ribosomal proteins quantify and contextualize genome recovery rates from metagenomes

Thanks to its diverse physiological properties that promote a variety of chemical gradients and surfaces (Bowen et al. 2018), the human oral cavity is home to diverse communities of microbes (Dewhirst et al. 2010). The human oral microbiome is a relatively well-characterized environment with a wealth of isolate genomes accessible through the Human Oral Microbiome Database (HOMD) (Escapa et al. 2018; T. Chen et al. 2010), and numerous genome-resolved metagenomics surveys that have captured representative genomes of microbial clades that have largely eluded cultivation efforts. Using EcoPhylo we first focused on a genome-resolved metagenomics survey which reconstructed multiple high-quality MAGs from tongue and plaque samples from the human oral cavity (Shaiber et al. 2020). While Shaiber et al. (2020) reported numerous genomes for elusive taxa, such as *Saccharimonadia* (TM7), *Absconditabacteria* (SR1), and *Gracilibacteria* (GN02), the genome-resolved metagenomic workflow failed to reconstruct MAGs that resolved to some of the best-represented organisms in culture collections from the oral cavity, such as members of the genus *Streptococcus*, (Escapa et al. 2018), which was represented by only two MAGs in Shaiber et al. (2020). This discrepancy compelled us to combine isolate genomes from the HOMD together with metagenomes and MAGs from Shaiber et al. (2020), to investigate whether EcoPhylo could reveal the differential recovery of genomes through distinct recovery approaches.

We started our analysis by combining 790 non-redundant MAGs and 14 metagenomic co-assemblies of tongue and plaque metagenomes reported by Shaiber et al (2020) with 8,615 isolate genomes we obtained from the HOMD (Supplementary Table 1). To characterize these data, we elected to use EcoPhylo with *rpL19* HMM, since it was the most frequent ribosomal protein with an average length of 393 nucleotides across all genomes in our collection, occurring in 98.59% of the HOMD genomes and 81.81% of the Shaiber et al. MAGs (Supplementary information). To assess the generalizability of observations made from *rpL19*, we also ran EcoPhylo on the same dataset with *rpS15* and *rpS2*, with the average length of 275 and 781 nucleotides, respectively (Supplementary Table 2, Supplementary information).

The EcoPhylo analysis of the *rpL19* genes found in the genomes and metagenomic assemblies resulted in a phylogenetic tree with 277 non-redundant bacterial representative sequences (Figure 2A, Supplementary Table 3). Hierarchical clustering of metagenomes based on the detection patterns of these *rpL19* sequences organized metagenomes into tongue and plaque sampling sites *de novo* (Figure 2A, Supplementary Figure 1), demonstrating that a single ribosomal gene family is able to capture the known ecological differences between these habitats. Many closely related *rpL19* genes that resolved to prevalent oral taxa, such as *Prevotella* and *Steptococcus*, showed within-genus differences in site specificity, a previously observed phenomenon (Eren et al. 2014) that is attributed to divergent accessory genomes (Mark Welch, Dewhirst, and Borisy 2019; Utter et al. 2020). Multiple ribosomal protein representative sequences recruited reads from tongue as well as plaque metagenomes, also matching prior observations of cosmopolitan taxa (Figure 2A, Supplementary information). Overall, the ecological insights revealed by *rpL19* recapitulated known ecology of oral microbes (Mark Welch, Dewhirst, and Borisy 2019) and provided a framework to assess genome recovery rates.

**Figure 2:**
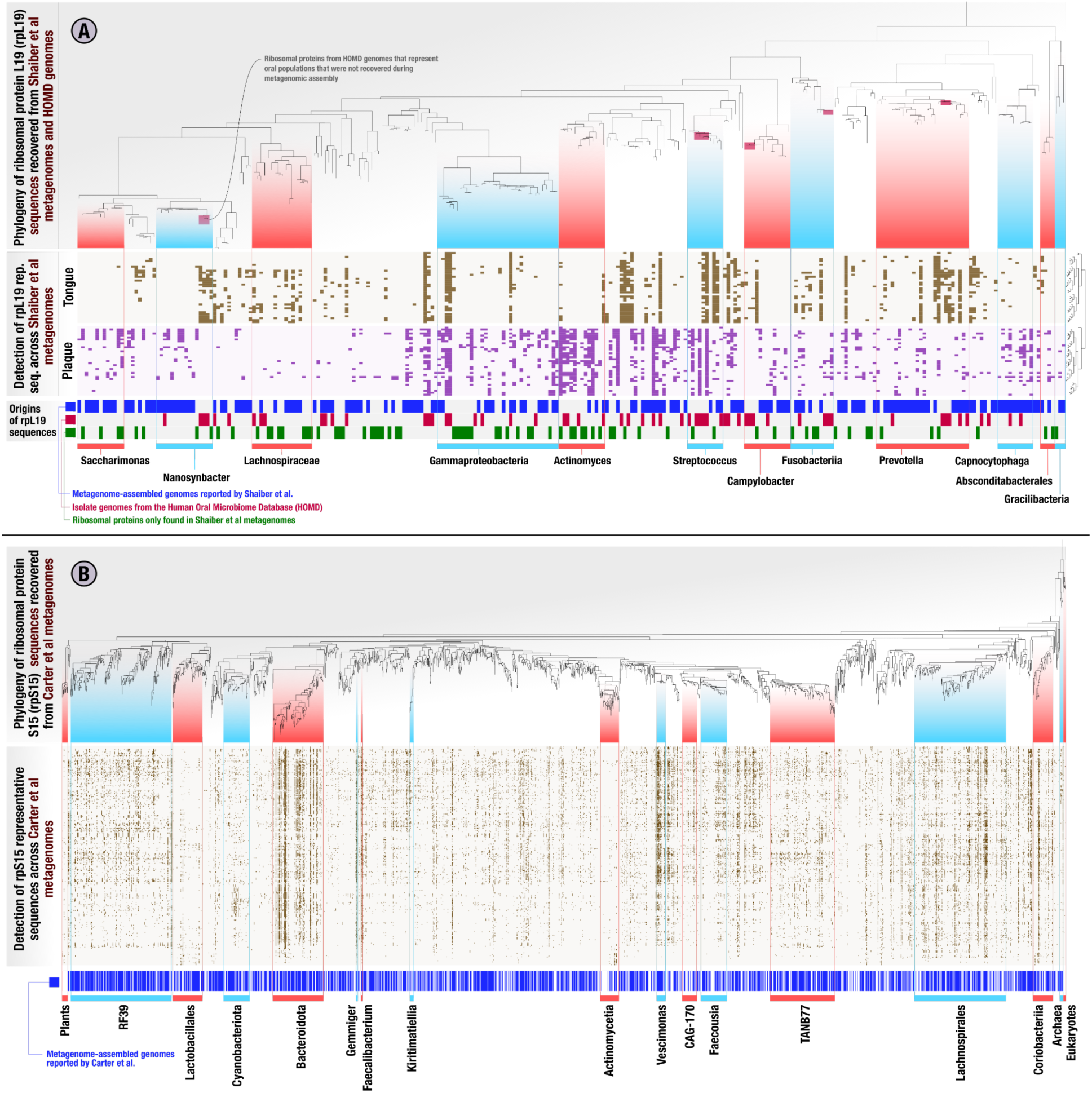
Ribosomal protein phylogeny and detection patterns across metagenomes from the human oral cavity and gut microbiomes. In the heatmaps in both panels, each column represents a ribosomal protein representative sequence, each row represents a metagenome, and each data point indicates whether a given ribosomal protein was detected in a given metagenome. The columns of heatmaps are ordered by a tree which represents a phylogenetic analysis of all ribosomal protein representative sequences, and the rows are ordered by a hierarchical clustering dendrogram that is calculated based on the ribosomal protein detection patterns across metagenomes. The panel **(A)** represents the EcoPhylo analysis of *rpL19* sequences across Shaiber et al. (2020) metagenome-assembled genomes (MAGs), Shaiber et al. (2020) oral metagenomes, and HOMD genomes, and includes three additional rows that indicate the origin of a given ribosomal protein, whether it is a metagenome-assembled genome (MAG, blue), HOMD isolate genome (red), or only recovered from metagenomic assemblies with no representation in genomes (green). Smaller red boxes in the phylogenetic tree mark microbial clades that were absent in the collection of MAGs and assemblies reported by Shaiber et al. (2020), but detected in Shaiber et al. (2020) metagenomes solely due to the inclusion of HOMD isolate genomes. The panel **(B)** represents the EcoPhylo analysis of *rpS15* sequences across the Carter et al. (2023) metagenome-assembled genomes (MAGs) and Carter et al. (2023) gut metagenomes from a Hadza tribe, and includes an additional row that indicates whether a MAG was reported for a given ribosomal protein (blue).

EcoPhylo tracks the origins of each sequence in each sequence cluster. Some *rpL19* clusters, representatives of which are shown in the phylogenetic tree in Figure 2A, were composed of sequences found only in metagenomic assemblies and not in MAGs or isolate genomes, highlighting clades present in the environment but not in genome collections. Other *rpL19* clusters only contained sequences represented in HOMD isolate genomes; despite their consistent detection in oral samples through metagenomic read recruitment, they were absent in metagenomic assemblies or MAGs, highlighting clades that are less accessible to short-read metagenomic assembly approaches (Figure 2). To calculate genome recovery rates for any given taxon, we divided the number of sequence clusters that contained a sequence from a given genome recovery method by the total number of representative sequences EcoPhylo reported for that taxon (Materials and Methods). This analysis revealed that 60.3% of the bacterial populations defined by *rpL19* gene clusters that were detected in metagenomic reads also appeared in MAGs. In other words, the overall bacterial MAG recovery rate in the study by Shaiber et al. (2020) was 60.3% (*rpS15*: 62.8%, *rpS2*: 53.2%) (Figure 2A, Supplementary Table 3, Supplementary Table 4). However, this rate of recovery was not uniform across individual taxa. EcoPhylo revealed higher MAG recovery rates for taxa such as *Saccharimonas* at 69.2% (*rpS15*: N/A, *rpS2:* 63.6%), and *Prevotella* at 76.9% (*rpS15*: 82.6%, *rpS2*: 84%). In contrast, the MAG recovery was lower for populations in other clades, including *Gammaproteobacteria* and *Fusobacteriia*, with MAG recovery rates of 47.1% (*rpS15*: 58.1%, *rpS2*: 44.8%) and 41.7% (*rpS15*: 46.2%, *rpS2*: 31.2%), respectively (Figure 2A, Supplementary Table 3, Supplementary Table 4). The MAG recovery rate was particularly low for *Streptococcus* at 30% (*rpS15*: 15.4%, *rpS2*: 10%), consistent with the presence of only two MAGs in Shaiber et al. (2020). However, the MAG recovery rate for *Actinomyces* was also very low at 23.1% (*rpS15*: 36.4%, *rpS2*: 13.3%) despite the characterization of nine *Actinomyces* MAGs by Shaiber et al. (2020) reveals a large number of distinct *Actinomyces* populations missed by MAGs even though they were present in the assemblies (Figure 2A, Supplementary Table 3, Supplementary Table 4). Overall, this analysis not only confirmed that MAG recovery rates are not uniform across microbial clades, but also showed that quantification of these rates is possible and may yield unexpected insights into the extent of diversity that is not represented in the final set of MAGs for some clades.

The inclusion of genomes from the HOMD increased the number of *rpL19* sequence clusters that contained genomes in this dataset, i.e., the total genome recovery rate, from 60.3% to 73.3% (*rpS15*: 74.8%, *rpS2*: 81.3%), and led to the representation of 35 additional microbial clades for which the metagenomic sequencing and analysis workflow implemented in Shaiber et al. (2020) did not assemble. As with MAGs, the improved detection of taxa among HOMD genomes was not uniform across clades (Figure 2A). For example, HOMD genomes offered genomic context for five additional *Streptococcus* populations, increasing the genome recovery rate from 30% with MAGs only, up to 80% when including the HOMD collection. When taking into account both MAGs and isolate genomes, the overall genome recovery rate of Shaiber et al. (2020) from a human oral microbiome dataset determined by EcoPhylo was 73.3%, showing that ribosomal protein phylogeography is an effective means to quantify genome recovery statistics for individual studies. Conversely, EcoPhylo results showed that 26.7% of the individual clades that could be detected through the presence of *rpL19* sequences in assemblies of Shaiber et al. (2020) metagenomes lacked genomic representation in both Shaiber et al. (2020) MAGs and HOMD isolates (Figure 2A). Clades that were solely detected through their assembled yet not binned ribosomal proteins increased the detection of populations of *Lachnospiraceae*, *Actinomyces*, *Gammaproteobacteria*, and *Patescibacteria* (Figure 2A). As EcoPhylo clusters ribosomal proteins at 97% nucleotide similarity, a conservative threshold that underestimates biodiversity by often grouping genomes with gANI below 95% (Olm et al. 2020), unbinned populations have the potential to contain novel genomic diversity.

Next, we applied EcoPhylo to another genome-resolved metagenomics study that recently characterized the gut microbiome of a Hadza hunter-gatherer tribe with a deep sequencing effort by Carter et al. (2023), in which the authors reported nearly 50,000 redundant bacterial and archaeal MAGs from 338 metagenomes with an average of 76 million paired-end reads (Supplementary Table 1). EcoPhylo analysis of this dataset with *rpS15* with an average length of 276 nucleotides, along with *rpS16* and *rpL19*, with the average length of 297 and 370 nucleotides respectively (Supplementary Figure 2), revealed a relatively high bacterial MAG recovery rate of 67.7% (*rpS16*: 72.8%, *rpL19*: 69.5%) (Figure 2B, Supplementary Table 2). While there were some clades, such as *Actinomycetia*, for which the genome recovery rate was as low as 31.9% (*rpS16*: 32.4%, *rpL19*: 33.3%), the high MAG recovery rate was generally uniform across all major taxa, highlighting how deep metagenomic sequencing yield relatively higher genome recovery rates (Supplementary Table 5, Supplementary Table 6, Supplementary Information).

Through these analyses, we are able to demonstrate that the MAGs obtained by Carter et al. (2023) more comprehensively represents the populations captured by their metagenomic assemblies of the human gut compared to the MAGs obtained by Shaiber et al. (2020) given their metagenomic assemblies of the oral cavity (Figure 2B, Supplementary Table 5, Supplementary Table 6, Supplementary Information). The ability to make such a statement highlights the utility of EcoPhylo at providing quantitative insights into the efficacy of genome-resolved surveys independent of biomes while offering a phylogenetic and biogeographical context for the populations that were detected in the assemblies.

Overall, EcoPhylo results from the human oral cavity and human gut ecosystems show that our workflow can scale to large metagenomic surveys, combine genomes from multiple sources to compare distinct recovery strategies at the level of individual phylogenetic clades, and recapitulate known ecological patterns.

## Discussion

Our work illuminates the efficiencies of current genome recovery methods and their ability to sample genomes from various microbiomes. By leveraging phylogenetically informative marker genes detected in metagenomic assemblies, such as ribosomal proteins, that are absent from final genome collections, EcoPhylo provides a robust framework for benchmarking genome recovery rates across multiple genome acquisition methods and contextualizing the ecological and evolutionary of genome collections with naturally occurring microbial populations. Our study examined two microbiome projects that used multiple genome recovery strategies (MAGs and isolate genomes) to survey the human oral cavity and gut. Overall, we found that the EcoPhylo workflow can quantitatively measure genome recovery rates and analyze heterogeneous genome collections to assess the efficacy of distinct recovery methods at the level of individual phylogenetic clades. In the oral microbiome, we found clade-specific biases of genome recovery rates across MAGs and isolate genomes. For example, isolate genomes effectively sampled *Streptococcus* while MAGs efficiently sampled *Prevotella*. However, in the human gut, we observed that deep metagenomic sequencing and subsequent generation of MAGs yielded higher genome recovery rates. Previous studies have shown that human gut samples exhibit relatively little genomic novelty (D. Wu et al. 2025), further supporting the effectiveness of genome-resolved metagenomics for acquiring genomes in this biome. Across both case studies, EcoPhylo generated insights into multi-domain ribosomal protein phylogeography and provided a valuable interactive data visualization strategy to evaluate the underlying microbial ecology of metagenomic sequencing projects.

The *de novo* profiling of ribosomal proteins in metagenomic assemblies resembles reference-based taxonomic profiling of metagenomic short reads to predict relative abundances of taxa, an idea that is implemented in multiple tools that use marker genes, such as Kraken (Wood and Salzberg 2014), MIDAS (Nayfach et al. 2016), Bracken (Lu et al. 2017), mOTUs (Ruscheweyh et al. 2022), and MetaPhlAn (Manghi et al. 2023), or processed conserved marker gene windows, such as SingleM (Woodcroft et al. 2024). As these tools typically report distinct taxa and their relative abundances, they indeed can help assess genome recovery efforts through direct comparisons of taxon names they identify to the taxonomy of recovered genomes. However, the requirement of a database of reference genomes and/or marker genes, and the absence of a direct link between the genes in assemblies and taxon names reported in tables limit applications with additional downstream opportunities such as targeted genome recovery. In contrast, the flexibility of surveying any marker gene, including ribosomal proteins, across user-provided metagenomic assemblies *de novo* offers an alternative approach that directly connects genes of unrecovered taxa to assemblies and estimates the number of populations detected in metagenomes regardless of their phylogenetic novelty across diverse samples and conditions.

While the phylogeography of ribosomal proteins offers valuable insights into genome recovery, these genes have notable limitations. First, ribosomal proteins assembled from metagenomes only capture a fraction of the microbial diversity in a sample, restricting genome recovery metrics to taxa detectable by metagenomics. Next, rates of evolution as well as the likelihood to be recovered through metagenomic assembly will differ across ribosomal protein families, complicating direct quantitative comparisons between different ribosomal proteins and in some cases will require surveying multiple ribosomal proteins to ensure the generalizability of observations from a single ribosomal protein. Additionally, individual ribosomal protein trees will have less phylogenetic power compared to concatenated ribosomal protein trees or longer marker genes. Although this may lead to suboptimal organism phylogenetics, the efficient organization of ribosomal proteins yields informative insights into the diversity of clades within a sample. Furthermore, when working with incomplete genomes, such as MAGs or SAGs, a single ribosomal gene family will rarely be detected across the entire genome collection and thus only a subset of genomes will be contextualized per protein. Yet the inherent trade-offs of using incomplete genomes (x ≥ 50% and less than 10% contamination) highlight ongoing challenges in genome recovery, as stricter completeness thresholds would further reduce the number of genomes available for analysis.

The modular design and customizable parameters of EcoPhylo allows users to go beyond ribosomal proteins and leverage other gene families tailored for specific analyses which can improve phylogenetics and the detection of specific taxa. For example, RNApolA and RNApolB have been leveraged for phylogeny-guided binning leading to the discovery of missing branches in viral evolution (Gaïa et al. 2023). Furthermore, phylogeography of functional protein families can be leveraged as proxies for microbial metabolism, e.g. phylogeography of ABC transporters can aid in modeling cryptic fluxes of microbial metabolites (Schroer 2023). The EcoPhylo workflow provides a platform for future microbiome projects to benchmark their genome recovery rates upon release of genome collections. Ribosomal protein phylogeography in tandem with reporting read recruitment percentages to representative genome collections, provides comprehensive insights into genome recovery rates given the biodiversity detected in metagenomes. The identification of microbial diversity hotspots with novel genomes using multi-biome databases is a growing trend in the field (Woodcroft et al. 2024; D. Wu et al. 2025). While these approaches enable broad-scale insights into habitats and samples with low genome recovery, EcoPhylo provides a direct link to the assembled genes within contigs of taxa with unrecovered genomes and illuminates phylogeny-guided genome recovery on a project-level. This immediate traceability to the underlying genomic fragments in metagenomic assemblies can catalyze targetted genome recovery. Future studies can leverage the strategy implemented in EcoPhylo to reanalyze existing metagenomic assemblies to identify missing clades or develop tailored methods to optimize overall genome recovery efforts by taking advantage of the increasing availability of genomes and metagenomes.

## Materials and Methods

### The EcoPhylo workflow

EcoPhylo is a computational workflow implemented in the open-source software ecosystem anvi’o (Eren et al. 2015, 2021) using the Python programming language and the workflow management system, Snakemake (Köster and Rahmann 2012). The primary purpose of EcoPhylo is to offer an integrated means to study phylogenetic relationships and ecological distribution patterns of sequences that match to any gene family based on user-provided hidden Markov model (HMM) searches from genomic and metagenomic assemblies. A minimal command line instruction to start an EcoPhylo run is ‘anvi-run-workflow -w ecophylo -c config.json’, where ‘anvi-run-workflow’ is a program in anvi’o that runs various workflows, and ‘config.json’ is a JSON formatted configuration file that describes file paths (such as the locations of genomes and/or metagenomes) and other parameters (such as the HMM to be used for a homology search, and sequence identity cutoffs). Comprehensive user documentation for EcoPhylo is available at https://anvio.org/m/ecophylo.

The minimum input for the EcoPhylo is a gene family hidden Markov model (HMM) and a dataset of genomic and/or metagenomic assemblies. EcoPhylo identifies and clusters target genes or translated proteins across assemblies to yield a non-redundant, representative set of open reading frames (ORFs). Next, an amino acid phylogenetic tree is calculated with the translated representative ORFs yielding the evolutionary history captured by homologues from input assemblies. An additional user input to the workflow is a metagenomic sequencing dataset representing ecological sampling or an experimental setup. With this input, the workflow performs metagenomic read recruitment against the representative ORFs to yield ecological insights into the gene family. Finally, the separate data types are integrated into a phylogeographic representation of the gene family (Figure 1).

The resulting sequences from the workflow can be organized in the EcoPhylo interactive interface either using an amino acid phylogenetic tree or using hierarchical clustering based on differential read recruitment coverage across metagenomic samples. Additionally, metagenomes can be hierarchically clustered based on the detection of the target gene family. It is recommended to employ hierarchical clustering of metagenomes or sequences in the EcoPhylo interactive interface with the detection read recruitment statistic (rather than coverage values) to minimize the effect of non-specific read recruitment (https://merenlab.org/anvio-views/).

An application of the EcoPhylo workflow with default settings will (1) identify gene families with the program ‘hmmsearch’ in (Eddy 2011) using the user-provided HMM model, (2) annotate affiliate hmm-hits with taxonomic names with ‘anvi-run-scg-taxonomy’ when applicable, (3) remove hmm-hits with less than 80% HMM model alignment coverage and incomplete ORFs with the anvi’o program ‘anvi-script-filter-hmm-hits’ with parameters ‘--min-model-coverage 0.8’ and ‘--filter-out-partial-gene-calls’ to minimize the inclusion of non-target sequences and spurious HMM hits, (4) dereplicate the resulting DNA sequences at 97% gANI and pick cluster representatives using MMseqs2 (Steinegger and Söding 2017), (5) use the translated representative sequences to calculate a multiple sequence alignment (MSA) using (Edgar 2004) with the ‘-maxiters 2’ flag (Edgar 2004), trim the alignment by removing columns of the alignment with trimAL with the ‘-gappyout’ flag (Capella-Gutiérrez, Silla-Martínez, and Gabaldón 2009), (6) remove sequences that have more than 50% gaps using the anvi’o program ‘anvi-script-reformat-fastà, (7) calculate a phylogenetic tree using FastTree (Price, Dehal, and Arkin 2010) with the flag ‘-fastest’, (8) perform metagenomic read recruitment analysis and profiling of non-translated representative sequences using the anvi’o metagenomic workflow (Shaiber et al. 2020), which by default relies upon Bowtie2 (Langmead and Salzberg 2012), (9) generate miscellaneous data to annotate the representative sequences including taxonomy with ‘anvi-estimate-scg-taxonomy’ for ribosomal proteins, cluster size, and sequence length, and finally (10) generate anvi’o artifacts that give integrated access to the phylogenetic tree of all representative sequences and read recruitment results that can be visualized using the anvi’o interactive interface and/or further processed for specific downstream analyses using any popular data analysis environment such as R and/or Python.

The workflow that resulted in the recovery and characterization of ribosomal proteins in our manuscript used the following additional steps: (1) we removed input reference genomes that were not detected in at least one of the input metagenomes above a detection value of 0.9 with their ribosomal protein (we kept all MAGs originating from the samples themselves) to only visualize detected populations, (2) we manually curated the ribosomal protein tree when necessary to remove sequences that appeared to be chimeric and those that formed spurious long branches likely originating from metagenomic assembly artifacts, and/or mitochondrial or plastid genomes (Supplementary information) and recalculated new amino acid phylogenetic trees with curated sequences with ‘FastTreè or IQTREE with the parameters ‘-m WAG -B 1000’ (Minh et al. 2020) and imported the new trees using the program ‘anvi-import-items-order’, and finally, (3) we generated additional metadata using in-house Python or R scripts and imported additional metadata using the program ‘anvi-import-misc-datà to decorate trees or metagenomes.

### Benchmarking EcoPhylo workflow with ribosomal proteins using CAMI synthetic metagenomes

We validated the EcoPhylo workflow by benchmarking it against the CAMI synthetic metagenomes (Meyer et al. 2022) to identify nucleotide clustering thresholds of ribosomal gene families to limit non-specific read recruitment while maximizing taxonomic resolution. We applied the EcoPhylo workflow across the three CAMI biome synthetic genomic/metagenomic datasets (Marine, Plant-associated, and Strain-madness). As an initial step, we identified the top five most frequent ribosomal gene families that were detected in single-copy in the associated genomic collections for each synthetic metagenomic dataset. We then conducted a parameter grid search, spanning 95%-100% nucleotide similarity parameter grid search (Meyer et al. 2022). Next, we measured the amount of non-specific read recruitment in each EcoPhylo iteration, i.e. reads with equal mapping scores between their primary and secondary alignments (multi-mapped reads), with the following Samtools command: ‘samtools view $sample | grep XS:i | cut -f12-13 | sed ‘s/..:i://g’ | awk ‘$1==$2’ | wc -l’. The percentage of non-specific read recruitment was calculated by dividing the number of multi-mapped reads by the total number of reads mapped to the representative dataset. With this, we identified that nucleotide clustering thresholds greater than 97% began to show signs of non-specific read recruitment (Supplementary information).

After identifying 97% nucleotide identity as the optimal threshold, we measured EcoPhylo’s ability to contextualize a genomic collection within metagenomic assemblies by quantifying the amount of genomic ribosomal genes clustering with their associated metagenomic assembly ribosomal gene (Supplementary information). Finally, we benchmarked the Shannon diversity and richness captured by different SCGs within the metagenomes and compared it to other taxonomic profiling tools submitted to CAMI (Meyer et al. 2022). To calculate Shannon diversity and richness values for SCGs processed by EcoPhylo we used the R package vegan (Dixon 2003) and Phyloseq (McMurdie and Holmes 2013). To calculate the richness and alpha diversity values for CAMI ground truth and other profiling tools we extracted relative abundance for each genera included in the associated biome files made available from CAMI. Shannon diversity for SCGs in the EcoPhylo were calculated with the anvi’o coverage statistic: Q2Q3 coverage. Datasets were cleaned and visualized with R packages in Tidyverse (Wickham et al. 2019).

### Genome collections

All MAG and SAG datasets were filtered for genomes with 50% completion and 10% redundancy using the single-copy core gene collections in anvi’o to meet medium-quality draft status in accordance with the community standards (Bowers et al. 2017). For the human oral cavity analysis, 8,615 human oral isolate genomes were downloaded from HOMD v10.1 (https://www.homd.org/ftp/genomes/NCBI/V10.1/) (Escapa et al. 2018) and 790 MAGs were downloaded from Shaiber et al. (2020) via (doi:10.6084/m9.figshare.12217805, doi:10.6084/m9.figshare.12217961).

For the Hadza tribe human gut microbiome analysis we followed the data download guidelines shared by Carter et al. (2023) to obtain genomes from doi:10.5281/zenodo.7782708. Carter et al. (2023) formed clusters at 95% gANI by including additional genomes outside of the MAGs they have reconstructed from the Hazda gut metagenomes. To exclusively analyze microbial genomes affiliated with the Hadza metagenomes, we filtered for cluster representatives with cluster members that contained at least one Hadzda adult or infant MAG which produced 2,437 representative Bacterial and Archaeal MAGs.

### Metagenome and metagenomic assembly datasets

To explore the phylogeography of ribosomal proteins, we used used 71 tooth and plaque metagenomes from the human oral cavity which were downloaded from the NCBI BioProject PRJNA625082 (Shaiber et al. 2020) along with associated co-assemblies (doi:10.6084/m9.figshare.12217799). Next, to explore deep sequencing in the human gut microbiome we used 388 metagenomes and assemblies from infant and adult members of the Hadza tribe (doi:10.5281/zenodo.7782708) using the FTP links shared in from the file ‘Supplemental_Table_S1.csv’ and NCBI BioProject PRJEB49206 (Carter et al. 2023). All metagenomes and associated assembly accessions can be found at Supplementary Table 1.

### Preprocessing of genomic and metagenomic assemblies and metagenomic short reads

Metagenomic and genomic assemblies were preprocessed with the anvi’o contigs workflow with the program ‘anvi-run-workflow -w contigs’ to predict open-reading frames with Prodigal (V2.6.3) and identify SCGs for taxonomic inference with ‘anvi-run-scg-taxonomy’ (Hyatt et al. 2010; Shaiber et al. 2020). No contig size filters were implemented during this process to include ribosomal proteins located on small contigs. To limit detection of misassemblies in downstream analyses, only ribosomal proteins with complete open-reading frames (as predicted by Prodigal) were analyzed with EcoPhylo (Hyatt et al. 2010). Additionally, metagenomic samples were quality controlled with the anvi’o metagenomics workflow with the program ‘anvi-run-workflow -w metagenomics’ (Shaiber et al. 2020). This workflow uses the tool ‘iu-filter-quality-minochè (Eren et al. 2013), which implements methods described in (Minoche, Dohm, and Himmelbauer 2011). All Snakemake workflows in this manuscript leveraged Snakemake v7.32.4 (Köster and Rahmann 2012).

### Gene-level taxonomy of ribosomal proteins

To assign gene level taxonomy to ribosomal proteins, the EcoPhylo workflow relies upon the anvi’o tools ‘anvi-run-scg-taxonomy’ and ‘anvi-estimate-scg-taxonomy’, which leverage the genomes and their taxonomy made available by the GTDB (Parks et al. 2022) to identify taxonomic affiliations of genes that match to any of the ribosomal proteins L1, L13, L14, L16, L17, L19, L2, L20, L21p, L22, L27A, L3, L4, L5, S11, S15, S16, S2, S6, S7, S8, or S9. During the workflow, EcoPhylo uses ‘anvi-run-scg-taxonomy’ to search for ribosomal genes annotated within each anvi’o contigs database against the downloaded marker gene dataset with DIAMOND v0.9.14 (Buchfink, Reuter, and Drost 2021). Later in the workflow, EcoPhylo runs ‘anvi-estimate-scg-taxonomy --metagenome-modè on the representative set of ribosomal proteins, which assigns a consensus taxonomy to each sequence. The program ‘anvi-estimate-scg-taxonomy’ does not provide a taxonomic annotation if the ribosomal protein is less than 90% similar to any of the ribosomal proteins found in GTDB genomes. In some cases, ribosomal proteins without taxonomic annotation can be manually annotated with taxonomy based on the annotated sequences that surround them in the phylogenetic tree, as we described in the section “Taxonomic binning to improve genome recovery estimations”.

### Selection of ribosomal proteins to contextualize genomic collections in metagenomes

To pick ribosomal gene families to study genome collections, we selected ribosomal genes that were annotated in the majority of genomes in single-copy. We then cross-referenced selected ribosomal genes with their assembly rates in metagenomes and disregarded candidate ribosomal gene families that were under- or over-assembled in the dataset. To do this, we ran the EcoPhylo workflow with the input dataset of genomic and metagenomic assemblies until the rule ‘process_hmm_hits’, which will filter for high-quality HMM-hits as described above. Finally, we extracted ribosomal protein hits from all assemblies with the anvi’o command ‘anvi-script-gen-hmm-hits-matrix-across-genomes’ and tabulated/visualized the distribution in R using the Tidyverse (Wickham et al. 2019).

### Distribution of HMM alignment coverage and SCG detection across GTDB

To identify optimal ribosomal proteins and HMM hit filtering thresholds, we explored the distribution of SCG detection and HMM alignment coverage across GTDB genomes. The analysis used the first two rules of the EcoPhylo workflow (anvi_run_hmms_hmmsearch and filter_hmm_hits_by_model_coverage) to annotate the RefSeq representative genomes from GTDB release 95 (Parks et al. 2020), with the single-copy core gene HMM collections included in anvi’o. The first rule of the workflow used the program ‘hmmsearch’ in (Eddy 2011)identify HMM hits, while the second rule was modified to include all HMM hit model coverage values by setting the parameter ‘anvi-script-filter-hmm-hits-table --min-model-coverage 0’. We stopped the workflow after this rule and visualized the raw distribution of model and gene coverage values from ‘hmmsearch --domtblout’ output file leading us to to identify an 80% HMM hit model coverage as an optimal filtering threshold to identify ribosomal proteins. Next, we restarted the workflow but re-modified the second rule parameter ‘anvi-script-filter-hmm-hits-table --min-model-coverage 0.8’ to filter for HMMs hits with at least 80% model alignment coverage. Finally, we extracted all ribosomal gene families from the genome dataset with anvi’o program ‘anvi-script-gen-hmm-hits-matrix-across-genomes’ and visualized the genome detection and SCG copy number across the dataset in in R using the Tidyverse (Wickham et al. 2019).

### Detection of whole genomes in metagenomic data

In some cases, ribosomal proteins clustering at 97% brought together large groups of highly similar isolate genomes. To identify the specific genome that is detected in the metagenomic datasets, we re-clustered the target EcoPhylo protein at 98% to resolve sequence clusters and thus increase the number of representative sequences. We then used the whole genomes associated with the new, larger set of representative proteins to explore their distribution in metagenomes by performing the anvi’o metagenomic workflow (Shaiber et al. 2020). Our threshold for detection of a whole-genome in metagenomic data was 50% (percent of genome covered by at least one read from metagenomic read recruitment), which was found to be efficient for human oral cavity microbes (Utter et al. 2020).

### Genome recovery rate estimations

Genome recovery rates were estimated to measure which individual or combination of genome types (MAGs, SAGs, isolate genomes) most effectively sampled clades in the ribosomal protein phylogenetic trees calculated during the EcoPhylo workflow. To calculate genome recovery rates for any given taxon, we divided the number of sequence clusters that contained a sequence from a given genome recovery method to the total number of representative sequences EcoPhylo reported for that taxon. Taxonomic assignments of sequence cluster representatives were determined with ‘anvi-estimate-scg-taxonomy’.

### Taxonomic binning to improve genome recovery estimations

A subset of ribosomal proteins lacked taxon assignments from ‘anvi-estimate-scg-taxonomy’ due to their sequence similarity being x < 90% to GTDB genomes (See methods section: Gene-level taxonomy of ribosomal proteins). Using the ‘anvi-interactivè interface, we examined the placement of these proteins in the EcoPhylo ribosomal protein phylogenetic tree and manually assigned taxon names based on the taxonomic affiliations of neighboring sequences. Unannotated sequences were assigned taxonomy only when phylogenetic clustering demonstrated clear consistency among neighboring sequences. These refined taxonomic annotations were used to improve estimations of genome recovery in the main figures (*rpL19* and *rpS15* in Figure 2).

## Data and code availability

The URL https://merenlab.org/data/ecophylo-ribosomal-proteins/ serves all code and data needed to reproduce our study. Additionally, all anvi’o artifacts that give interactive access to EcoPhylo interfaces are publicly available at doi:10.6084/m9.figshare.28207481. Publicly available genomes and metagenomes we used in our study are listed in the Supplementary Tables. Supplementary Tables as well as Supplementary Information text are available at doi:10.6084/m9.figshare.28847705.

## Supplementary Information

doi:10.6084/m9.figshare.28847705

## Acknowledgements

We thank all authors who made the raw metagenomic reads, metagenomic assemblies, and genomes from their manuscripts publically available and conveniently accessible for secondary analyses. We also thank the members of the Meren Lab (https://merenlab.org/people/), the Light Lab (https://www.lightlab.uchicago.edu/people/), and the Blekhman Lab (http://blekhmanlab.org/members.html) for helpful discussions and whiteboard sessions. We are also thankful to Pedram Esfahani, Kimberly Grasch, and the rest of the University of Chicago Center for Research and Computing Center for their patience and support. MSS acknowledges support from NIH Genetics and Regulation Training Grant (T32 GM07197). AME acknowledges support from the Center for Chemical Currencies of a Microbial Planet (C-CoMP) (NSF Award OCE-2019589, C-CoMP publication #070), and Simons Foundation (grant #687269).

## Author contributions

MSS and AME conceptualized the study. MSS curated data and performed formal analyses. MSS, IAV, MLK, MS, SEM, and AME developed software tools. MSS, FT, and AME interpreted findings. LM, TOD, and SHL helped with interpretation of results. MSS and AME wrote the original draft of the study. SHL and AME managed the project and acquired funding. All authors commented on and made suggestions, and approved the final manuscript.

